# Inactivation of Highly Transmissible Livestock and Avian Viruses Including Influenza A and Newcastle Disease Virus for Molecular Diagnostics

**DOI:** 10.1101/2023.09.13.557451

**Authors:** Jennifer L. Welch, Ram Shrestha, Heather Hutchings, Narinder Pal, Randall Levings, Suelee Robbe-Austerman, Rachel Palinski, Karthik K. Shanmuganatham

## Abstract

There is a critical need for an inactivation method that completely inactivates pathogens at the time of sample collection but maintains the nucleic acid quality required for diagnostic PCR testing. This inactivation method is needed to alleviate concerns about transmission potential, reduce shipping complications and cost, and allow testing in lower containment laboratories to improve disease diagnostics by improving turn-around time. This study evaluated a panel of ten surrogate viruses that represent highly pathogenic animal diseases. These results showed that a commercial (PrimeStore®) molecular transport media (PSMTM) completely inactivated all viruses tested by >99.99% as determined by infectivity and serial passage assays. However, detection of viral nucleic acid by qRT-PCR was comparable in PSMTM and control-treated conditions. These results were consistent when viruses were evaluated in the presence of biological material such as sera and cloacal swabs to mimic diagnostic sample conditions for non-avian and avian viruses, respectively. The results of this study may be utilized by diagnostic testing laboratories for highly pathogenic agents affecting animal and human populations. These results may be used to revise guidance for select agent diagnostic testing and shipment of infectious substances.

**Contribution to the field:** Active surveillance and confirmatory testing efforts are in place to protect animals in the United States from certain highly contagious diseases and to limit financial impacts to consumers and producers when the food supply is disrupted. Confirmatory testing typically utilizes nucleic acid detection to identify active infection. Testing is required to be completed in high containment facilities due to the elevated pathogenicity and impact potential of animal diseases. The requirement for testing in high containment facilities limits the ability for regional and state laboratories to test for Tier 1 select agents. Shipment of diagnostic samples is costly, as well as time and temperature sensitive to avoid deterioration of sample quality needed for testing. These constraints lengthen response time and testing turn-around time. Here, we showed that a commercial (PrimeStore®) molecular transport media (PSMTM) completely inactivated all viruses tested without affecting nucleic acid detection/integrity. These data suggest that highly contagious agents are effectively inactivated by PSMTM without compromising the nucleic acid needed for diagnostic testing. These data provide support that this inactivation method can be utilized during sample collection to reduce constraints in disease diagnostics and in reagent sharing among international laboratories.

## Introduction

Prevention efforts are in place to protect animals in the United States from highly contagious diseases caused by classical swine fever virus (CSFV), African swine fever virus (ASFV), foot-and-mouth disease virus (FMDV), eastern equine encephalitis virus (EEEV), Newcastle disease virus (NDV), and highly pathogenic avian influenza A virus (HPAI) among others [1-3]. When an approved vaccine or treatment is unavailable, depopulation is the mandated course of action for affected farms [4]. This places critical importance on prevention efforts, as depopulation disrupts the food supply and has major financial implications for producers and consumers [1]. For example, the largest ASF outbreak to date was reported in China in 2018-2019 and is estimated at more than $111.2 billion in total financial losses [5]. The ongoing HPAI outbreak was first identified in the United States in 2022, and current US economic losses are estimated to range $2.5-3 billion [6]. In addition, some of these viruses including FMDV, EEEV, NDV, and HPAI are zoonotic thus are a concern for public health [7-10]. Prevention efforts include active surveillance testing [11].

Active surveillance testing requires shipment of suspect samples of highly contagious diseases to BSL3+ high containment facilities for PCR confirmatory testing to meet select agent requirements [12, 13]. Shipment of potentially infectious substances is costly, time-sensitive, requires specialized shipping containers, and requires a cold-chain to avoid deterioration of samples for PCR testing [14]. An inactivation method that protects the nucleic acid content needed for PCR testing while removing the threat of transmission and minimizing the temperature requirement is urgently needed to reduce shipping costs and derestrict confirmatory testing to lower biosafety-level facilities (BSL2). This will alleviate the strain the high PCR testing demand is placing on limited BSL3 facilities across the United States, thereby decreasing testing response time and improving practical difficulties. This includes improving the complications of sharing reagents among international laboratories [15]. The accidental shipment of live anthrax to laboratories within the United States due to incomplete inactivation illustrates the importance of an inactivation method that provides a safety buffer for use with high consequence pathogens [16].

PrimeStore® molecular transport media (PSMTM) tubes contain proprietary reagents that inactivate both viruses and bacteria by disrupting lipid membranes and inhibiting replication machinery while stabilizing nucleic acids (PrimeStore®, Longhorn Vaccines and Diagnostics)[17, 18]. As indicated by the publicly available safety data sheet, this reagent contains a mixture of guanidine thiocyanate, ethanol, and n-lauroylsarcosine [19]. Guanidine thiocyanate-based reagents have previously been shown to effectively inactivate poliovirus and FMDV, though comparison of results should be interpreted with caution as exact formulation of PSMTM is proprietary [20, 21]. Previous studies have shown that ethanol and n-lauroylsarcosine inactivate various enveloped and some nonenveloped viruses by interacting with lipids and denaturing proteins [22].

PSMTM is authorized by the Food and Drug Administration (FDA) as a Class II device for collection of samples suspected of containing influenza A virus (IAV) and *Mycobacterium tuberculosis* [17].

Other studies have shown that PSMTM effectively inactivates SARS-CoV-2, adenovirus type 5, influenza A H3N2, and HPAI H5N1 at ambient temperature [17, 23]. Influenza A H1N1 studies showed that viral RNA was preserved in PSMTM for 30 days at 25°C, and cycle threshold (CT) values were minimally reduced from day 0 to day 30 [23]. In addition, PSMTM tubes were utilized during the COVID-19 pandemic for SARS-CoV-2 clinical testing [17, 24]. Studies showed 10 minutes of contact time was sufficient to completely inactivate SARS-CoV-2, and no virus was detected in titering assays or after serial passage on susceptible cells [24]. Together, these data support that PSMTM inactivates viral pathogens and preserves nucleic acids at ambient temperature, thereby permitting normal shipping procedures without disrupting normal PCR testing workflows.

In this study, we assess the effectiveness of PSMTM at inactivation of surrogate viruses that represent USDA Veterinary Services (VS) select agent viruses, some of which are considered foreign animal disease agents including ASFV, CSFV, and FMDV (Table 1) [3]. Due to the high pathogenicity and transmission consequences of these viruses, it is important to validate PSMTM with diverse viruses, sample types, and collection conditions. Surrogate viruses were utilized in this study as inactivation testing was completed in BSL2 conditions. The selected surrogate viruses represent viruses with diverse classifications in exterior structure (envelope vs nonenveloped), nucleic acid composition (DNA vs RNA), and nucleic acid structure (single vs double strand) (Table 1). At least one representative virus was utilized for each target virus and selected based on: 1. ease of ability to grow to high titers; 2. accessibility; 3. relevance to target virus genus and family.

**Table 1:**
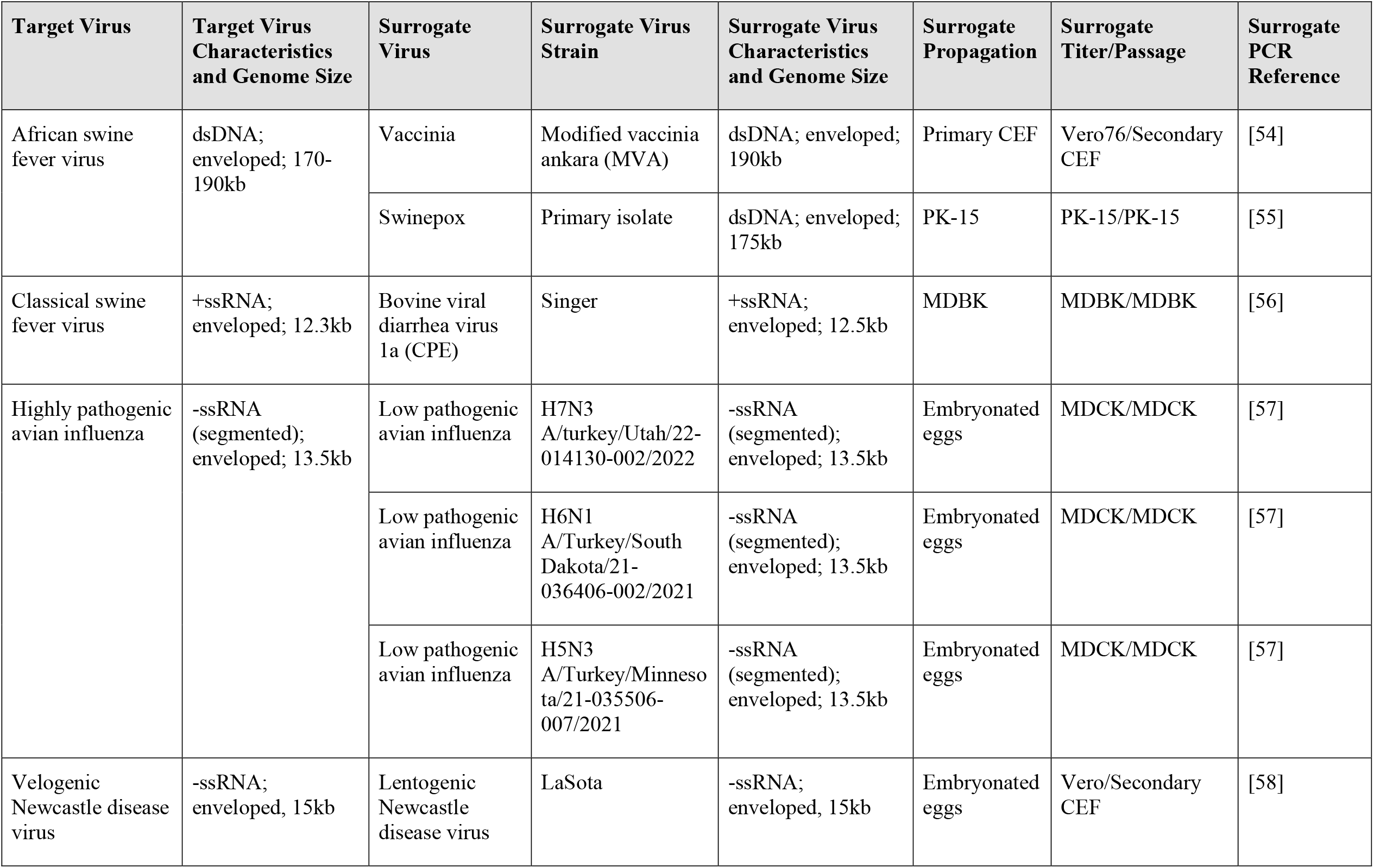
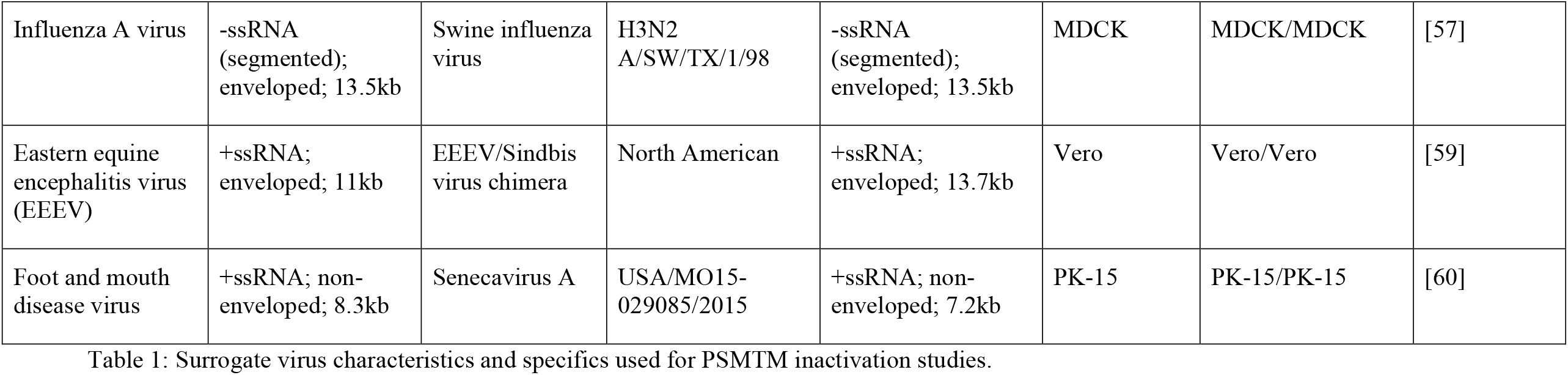
Surrogate virus characteristics and specifics used for PSMTM inactivation studies.

| Target Virus | Target Virus Characteristics and Genome Size | Surrogate Virus | Surrogate Virus Strain | Surrogate Virus Characteristics and Genome Size | Surrogate Propagation | Surrogate Titer/Passage | Surrogate PCR Reference |
| --- | --- | --- | --- | --- | --- | --- | --- |
| African swine fever virus | dsDNA; enveloped; 170-190kb | Vaccinia | Modified vaccinia ankara (MVA) | dsDNA; enveloped; 190kb | Primary CEF | Vero76/Secondary CEF | [54] |
|  |  | Swinepox | Primary isolate | dsDNA; enveloped; 175kb | PK-15 | PK-15/PK-15 | [55] |
| Classical swine fever virus | +ssRNA; enveloped; 12.3kb | Bovine viral diarrhea virus 1a (CPE) | Singer | +ssRNA; enveloped; 12.5kb | MDBK | MDBK/MDBK | [56] |
| Highly pathogenic avian influenza | -ssRNA (segmented); enveloped; 13.5kb | Low pathogenic avian influenza | H7N3 A/turkey/Utah/22-014130-002/2022 | -ssRNA (segmented); enveloped; 13.5kb | Embryonated eggs | MDCK/MDCK | [57] |
|  |  | Low pathogenic avian influenza | H6N1 A/Turkey/South Dakota/21-036406-002/2021 | -ssRNA (segmented); enveloped; 13.5kb | Embryonated eggs | MDCK/MDCK | [57] |
|  |  | Low pathogenic avian influenza | H5N3 A/Turkey/Minnesota/21-035506-007/2021 | -ssRNA (segmented); enveloped; 13.5kb | Embryonated eggs | MDCK/MDCK | [57] |
| Velogenic Newcastle disease virus | -ssRNA; enveloped, 15kb | Lentogenic Newcastle disease virus | LaSota | -ssRNA; enveloped, 15kb | Embryonated eggs | Vero/Secondary CEF | [58] |
| Influenza A virus | -ssRNA<br>(segmented);<br>enveloped; 13.5kb | Swine influenza<br>virus | H3N2<br>A/SW/TX/1/98 | -ssRNA<br>(segmented);<br>enveloped; 13.5kb | MDCK | MDCK/MDCK | [57] |
| Eastern equine<br>encephalitis virus<br>(EEEV) | +ssRNA;<br>enveloped; 11kb | EEEV/Sindbis<br>virus chimera | North American | +ssRNA;<br>enveloped; 13.7kb | Vero | Vero/Vero | [59] |
| Foot and mouth<br>disease virus | +ssRNA; non-<br>enveloped; 8.3kb | Senecavirus A | USA/MO15-029085/2015 | +ssRNA; non-<br>enveloped; 7.2kb | PK-15 | PK-15/PK-15 | [60] |

ASFV is the only member of the Asfarviridae family, making surrogate selection a challenge. To address this, we utilized species-relevant swinepox virus and the genetically similar virus, vaccinia virus [25-27]. Both swinepox and vaccinia virus are members of the Poxvirus family and like Asfarviridae, are nucleocytoplasmic large DNA viruses (NCLDV) that share a common ancestor [28]. Surrogate selections for other target viruses were previously identified by others and included bovine viral diarrhea virus (BVDV) as surrogate for CSFV and senecavirus A (SVA) as surrogate for FMDV [29, 30]. In addition, less pathogenic strains of target viruses were used as surrogates when possible, such as low pathogenic avian influenza (LPAI) H7N3, LPAI H6N1, and LPAI H5N3 as surrogates for HPAI, lentogenic NDV as surrogate for velogenic NDV, swine influenza virus (SIV) as surrogate for IAV, and EEEV/sindbis virus chimera as surrogate for EEEV [31]. Therefore, surrogate virus selections were best matched to the target viruses for NDV, HPAI, and IAV (same species, genus, and family) followed by EEEV (chimera of target virus and same genus), CSF (same genus and family), FMDV (same family), and ASF (common ancestor).

Diagnostic samples are collected in biological material that may influence inactivation efficacy. Previous studies identified that the presence of blood may interfere with viricidal activity of disinfectants [32]; however, other studies have shown that viruses remained sensitive to chemical inactivation in the presence of blood [33]. Here, we evaluate PCR detection and inactivation of virus replication after PSMTM treatment. In addition, inactivation was also evaluated for sera and cloacal swabs. The results of this study show PSMTM completely inactivated all viruses tested without compromising nucleic acid quality for PCR testing.

## Materials and methods

### Viruses

EEEV/sindbis virus chimera, swinepox, SVA, LPAI H6N1, LPAI H7N3, LPAI H5N3, SIV H3N2, lentogenic NDV, and BVDV 1a were provided by the USDA Diagnostic Virology Laboratory (DVL). Vaccinia virus was purchased from the American Type Culture Collection (ATCC). Viruses were propagated and titered by median tissue culture infectious dose (TCID_50_) in appropriate egg or cell systems (Table 1), and all eggs and cells used for propagation and titering were provided by USDA Diagnostic Bioanalytical and Reagents Laboratory (DBRL) Proficiency Testing and Reagents (PTR) section. Viruses were propagated in minimum essential media (MEM) with 5% fetal bovine serum, 4mM L-glutamine (Gibco), 0.015g/l penicillin, 0.1g/l streptomycin, and 1x non-essential amino acids (Gibco). Virus titrations were performed in MEM media with 2% fetal bovine serum and the same supplements as listed for the propagations.

### Virus natural decay inactivation

200uL of virus were added to one well of a 24 well tissue culture plate in triplicate for each timepoint. Viruses were incubated at ambient temperature until the indicated timepoint. To mitigate the effects of evaporation, viruses were collected at the indicated timepoint by addition of 200uL of virus propagation media to each well and repeated pipetting.

Collected virus was then frozen once at -80°C until titration by TCID_50_ on susceptible cells (Table 1).

### Virus PSMTM inactivation

Viruses were treated with PrimeStore® molecular transport media (PSMTM) according to manufacturer instructions. 1 part virus was combined with 3 parts PSMTM and incubated for 1 hour at ambient temperature. Virus infectivity (TCID_50_) was compared to no inactivation control where phosphate-buffered saline (PBS) replaced PSMTM and was treated the same. To mimic relevance to animal diagnostic samples collected in biological material, viruses were combined with species-appropriate serum or swab. For sera studies, virus was diluted 1:1 with swine or bovine serum prior to treatment with PSMTM or PBS. For swab studies, each virus in liquid media was added to a tube containing a cloacal swab collected from an uninfected chicken then incubated at ambient temperature ∼10 minutes. The virus in liquid media was then collected by pipette and treated with PSMTM or PBS control as described above. All control and inactivation procedures were completed in 3 experimental replicates. 3 parts PSMTM or PBS control to 1 part virus ratio was maintained for all experiments except for the varying ratio experiments. Ratio variations of 1 part PSMTM or PBS control to 1 part virus, 1 part PSMTM or PBS control to 3 parts virus, 1 part PSMTM or PBS control to 10 parts virus, and 1 part PSMTM or PBS control to 100 parts virus were also evaluated with EEEV/sindbis virus chimera to assess PSMTM inactivation if the recommended ratio is not maintained.

### Virus serial passage

PSMTM or PBS control treated viruses were passaged on susceptible cells (Table 1) for a minimum of 3 serial passages. PSMTM is highly cytotoxic to cells (SFig. 1). To minimize cytotoxicity of PSMTM, buffer exchange was completed prior to first passage for PSMTM and PBS control treated viruses. PSMTM and PBS control treated virus material were diluted 1:10 in MEM media without supplements. Diluted material was then added to an Amicon® 50kDa cutoff centrifugal filter (Millipore) and centrifuged at 1500 x *g* until diluted material was concentrated to the original volume (∼10 minutes). This filter cutoff was chosen as all viruses included in this study were larger than 50kDa and would be retained in the filter reservoir. Concentrated material was then diluted again 1:10 in MEM media and 50kDa concentration steps were repeated. After final concentration to the original volume, PSMTM and PBS treated virus were diluted 1:10 in MEM virus propagation media, and the total volume (2.0mL) was added to one well of a 6-well plate. Cells were incubated with PSMTM or PBS treated viruses until cytopathic effect (CPE) in PBS control treated virus wells or a maximum of 7 days. Passaged products were collected by three freeze/thaw cycles of cells and supernatant together. Passaged products were then centrifuged at 1500 x *g* for 10 minutes to remove cell debris. A fraction (10%) of the clarified product was diluted 1:10 in MEM virus propagation media, and the total volume (2.0mL) was added to naïve cells (one well of a 6-well plate) for each subsequent passage. Collection of PSMTM inactivated or control treated viruses from susceptible cells were designated as passaged virus (Fig. 2-4).

### Virus PCR detection

Nucleic acid was extracted from 200uL virus material treated with PSMTM or PBS and each passage product from serial passages described above. Nucleic acid was extracted using the MagMax™ CORE nucleic acid purification kit (Applied Biosystems) on KingFisher Flex (ThermoScientific) instrument with provided MagMax_CORE_Flex script (with heat) according to manufacturer instructions. Nucleic acid was detected by virus-specific primer/probe according to indicated reference sequences (Table 1) utilizing TaqMan™ Fast Virus 1-Step Master Mix (Applied Biosystems) and QuantStudio™ 5 (ThermoScientific) real-time PCR instrument. Cycling conditions for LPAI H6N1, LPAI H7N3, LPAI H5N3, SIV H3N2, NDV, and BVDV were as follows: 50°C for 5 min, 95°C for 20 s, followed by 45 cycles of 95°C for 15 s and 60°C for 45 s. Cycling conditions for vaccinia virus were as follows: 95°C for 20 s, followed by 45 cycles of 95°C for 10 s and 60°C for 30 s. Cycling conditions for EEEV/sindbis virus chimera and SVA were as follows: 50°C for 15 min, 95°C for 2 min, followed by 45 cycles of 95°C for 15 s and 60°C for 30 s. Cycling conditions for swinepox were as follows: 50°C for 5 min, 95°C for 20 s, followed by 45 cycles of 95°C for 15 s and 53°C for 1 min. Data were analyzed utilizing Design and Analysis software (version 2.6, ThermoScientific) where threshold values were set at 0.100. A CT of 45 was used as the no amplification negative threshold. Comparison of stock virus TCID_50_ and CT values revealed similar trends, validating assay parameters (SFig. 2).

### Virus PCR stability

200uL PSMTM or PBS control treated viruses were added to screw cap tubes in triplicate for each timepoint. Screw cap tubes were utilized to mimic PSMTM manufacturer tube conditions that may be utilized in field sample collection. Timepoints were collected for a maximum of 21 days to mimic the maximum anticipated transit time once a field sample is collected to when it reaches a laboratory for diagnostic testing. PSMTM or PBS control treated viruses were incubated at ambient temperature until the indicated timepoint where viruses were collected by addition of 200uL of virus propagation media to each tube and repeated pipetting. Collected virus was frozen once at -80°C until titration by TCID_50_ on susceptible cells (Table 1) and PCR detection performed as described above.

### Cell viability

Cell viability was determined by MTT (Invitrogen) assay where the cell lines utilized in this study were plated in 96-well plates and incubated with PSMTM or PBS control dilutions.

PSMTM or PBS were initially diluted to the manufacturer recommended ratio used for PSMTM inactivation studies described above where virus propagation media (not containing virus) was substituted for 1 part virus to assess the effect of PSMTM alone (without virus) on cell viability as these viruses induce CPE and would confound assessment of the effect of PSMTM on cell viability. PSMTM or PBS control treated media were then serially diluted 1:10 and incubated with cells for 48 hours. Cells were then incubated with 5 mg/mL MTT reagent in the dark for 3.5 hours at 37°C before addition of 4mM HCl in isopropanol and monolayer disruption by pipetting. Absorbance was read at 490nm using a BioTek plate reader. PSMTM cell viability values were normalized to equivalent PBS dilution set at 100%.

### Statistics

Statistical significance was determined by GraphPad software v9.3.1 (GraphPad Prism 9) using two-tailed Student’s t-test. *P* values < 0.05 were considered significant. Error bars represent standard error of the mean (SEM) from three biological replicates.

## Results

### Diverse viruses show similar patterns of stability at ambient temperature

To evaluate the potential risk of shipping transmissible high containment samples without an inactivation method applied at the time of sample collection, we evaluated the natural decay over time of the viruses used in this study. Virus infectivity at each timepoint was compared to timepoint The timepoint 0 condition shows that a minimum of 4-log virus was deposited and recovered for all viruses tested (Fig 1A-J).

**Figure 1:**
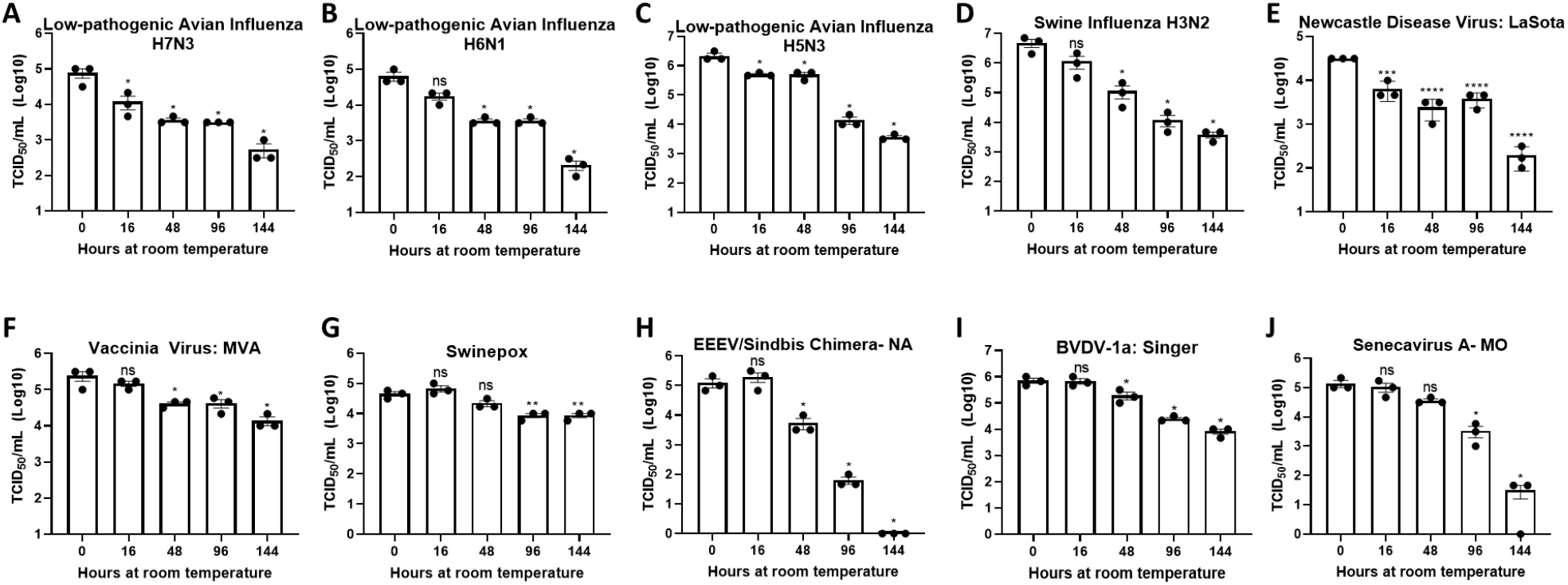
Diverse viruses are stable at ambient temperature. Virus recovery over time deposited on a plastic surface at ambient temperature for A) LPAI H7N3; B) LPAI H6N1; C) LPAI H5N3; D) SIV H3N2; E) NDV; F) Vaccinia; G) Swinepox; H) EEEV/Sindbis chimera; I) BVDV; and J) Senecavirus A. Virus recovery was assessed by TCID_50_ titer using the cell line described in Table 1. Each timepoint was compared to timepoint 0. Significance was determined using the Student’s t-test. * *p*<0.05; ** *p*<0.01; *** *p*<0.001; **** *p*<0.0001; ns, not significant. Error bars represent standard error of the mean (SEM) of triplicate experiments.

Although natural decay varied between individual viruses, results show a maximum reduction of 1-log after 48 hours at ambient temperature and survival of all viruses after 144 hours at ambient temperature with the exception of EEEV/sindbis virus chimera, which was completely inactivated at 144 hours (Fig 1A-J). These data indicate that highly pathogenic viruses may survive prolonged periods of time under natural conditions and demonstrate the need for an inactivation method that can be utilized at the time of sample collection to reduce the risk of transmission.

### PSMTM completely inactivates viruses representing highly contagious animal diseases

Viruses were treated with PSMTM according to manufacturer recommended conditions or PBS (no PSMTM) for control comparison. Viral replication was assessed by TCID_50_ and susceptible cell serial passage, and nucleic acid was evaluated by qRT-PCR. PSMTM treatment completely inactivated all viruses tested compared to PBS control. There was no infectivity remaining in PSMTM-treated viruses and virus titers were reduced >4-log (99.99%) (Fig. 1A; Table 2). Despite no remaining virus infectivity post PSMTM-treatment, CT values of viruses treated with PSMTM compared to no PSMTM control were highly comparable and varied by less than 1.2 CT value (Fig. 1B; Table 2). A CT value change of 3.3 is considered to represent a 1-log (10-fold) change [34]. To ensure complete inactivation, PSMTM- and control-treated viruses were serially passaged on susceptible cells (Table 1). CT values of PSMTM-treated viruses reached negative threshold with no observable cell infectivity by passage endpoint of a minimum of three passages, indicating no nucleic acid amplification or virus replication (Fig 2C; SFig3-4). Conversely, CT values of control-treated viruses were stable or showed reduced CT (increased nucleic acid) at passage endpoint compared to input (Fig 2C; SFig 3). Cell infectivity was also observed at passage endpoint for control-treated viruses (SFig 3). Together, these data show complete inactivation of PSMTM-treated virus replication without deterioration of the nucleic acid needed for PCR testing.

**Figure 2:**
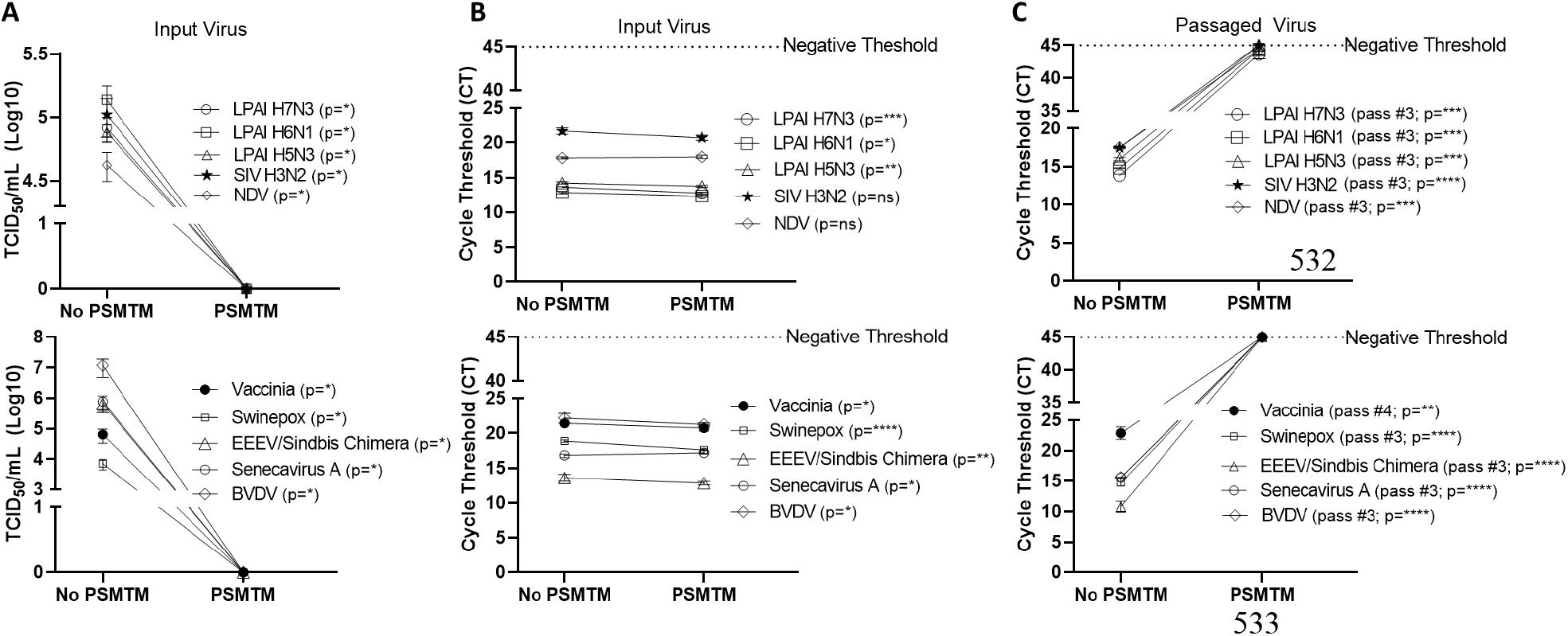
PSMTM effectively inactivates diverse animal viruses while maintaining detection of nucleic acid. A) Virus recovery after PSMTM inactivation or PBS no-inactivation control. Virus recovery was assessed by TCID_50_ titer using the cell line described in Table 1. B) Nucleic acid cycle threshold (CT) detection after PSMTM inactivation or PBS no-inactivation control. C) Serial passage endpoint nucleic acid cycle threshold (CT) detection after PSMTM inactivation or PBS no-inactivation. Serial passage was completed using the cell line in Table 1 and description outlined in methods. Nucleic acid was detected according to the reference in Table 1 and description in methods. PSMTM-treated viruses were compared to corresponding no-PSMTM control and inactivation was assessed using manufacturer recommended conditions. Significance was determined using the Student’s t-test. * *p*<0.05; ** *p*<0.01; *** *p*<0.001; **** *p*<0.0001; ns, not significant. Error bars represent standard error of the mean (SEM) of triplicate experiments.

**Table 2:**
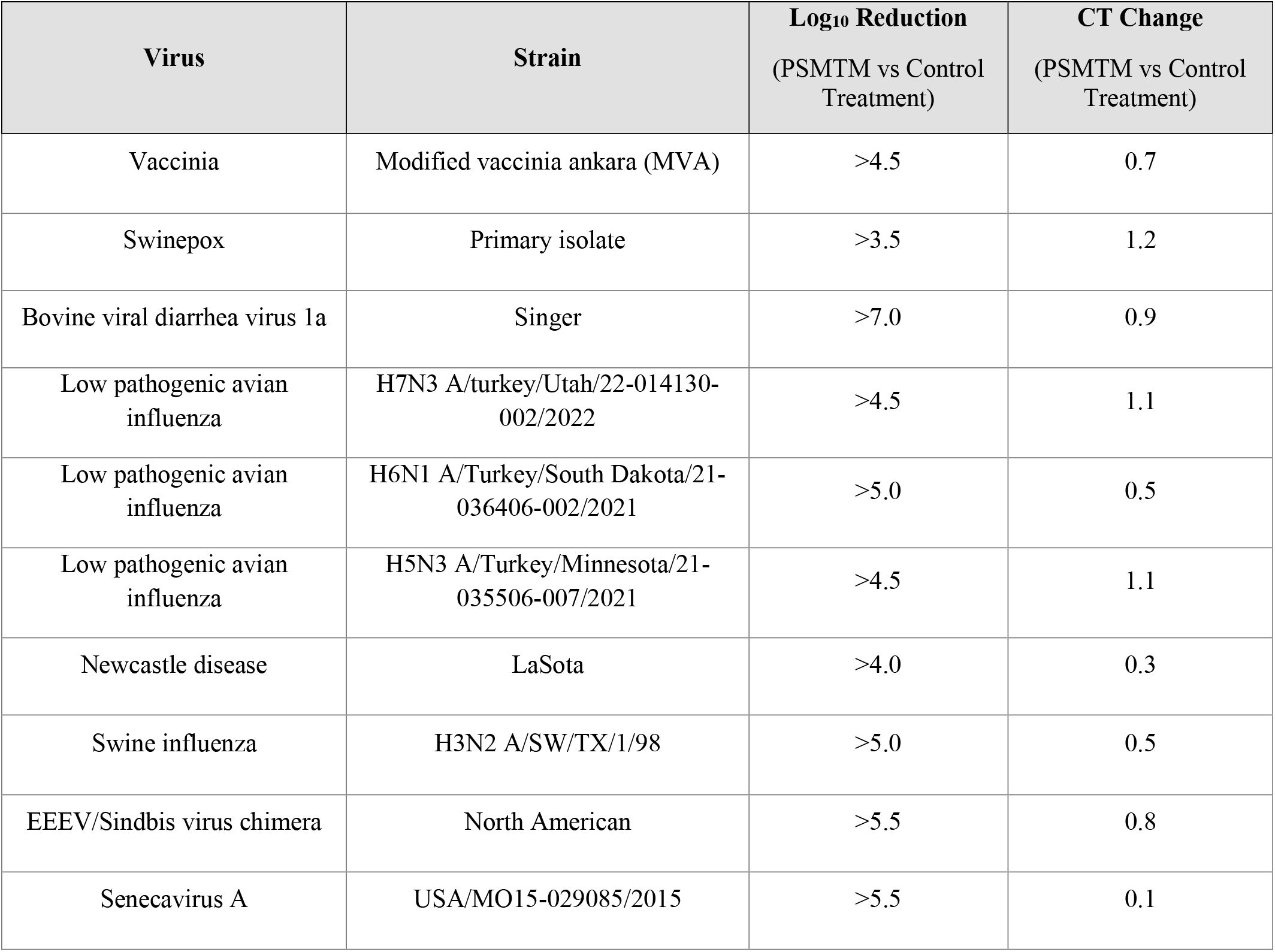
Log10 TCID_50_ reduction and qRT-PCR cycle threshold (CT) change of PSMTM treatment vs control treatment viruses.

We assessed PSMTM inactivation of viruses in the presence of species-relevant sera or, in the case of avian surrogates, cloacal swab. Consistent with our findings without added biological material, PSMTM treatment completely inactivated all virus-spiked sera or virus-spiked cloacal swabs (Fig. 3A). CT values of virus-spiked sera or virus-spiked cloacal swabs treated with PSMTM were comparable to no-PSMTM control (Fig. 3B). Importantly, serial passage of PSMTM-treated viruses in the presence of serum or a swab approached negative threshold by passage endpoint whereas control-treated viruses did not (Fig. 3C; SFig 3). These data show that the presence of biological material does not affect PSMTM inactivation of viruses.

**Figure 3:**
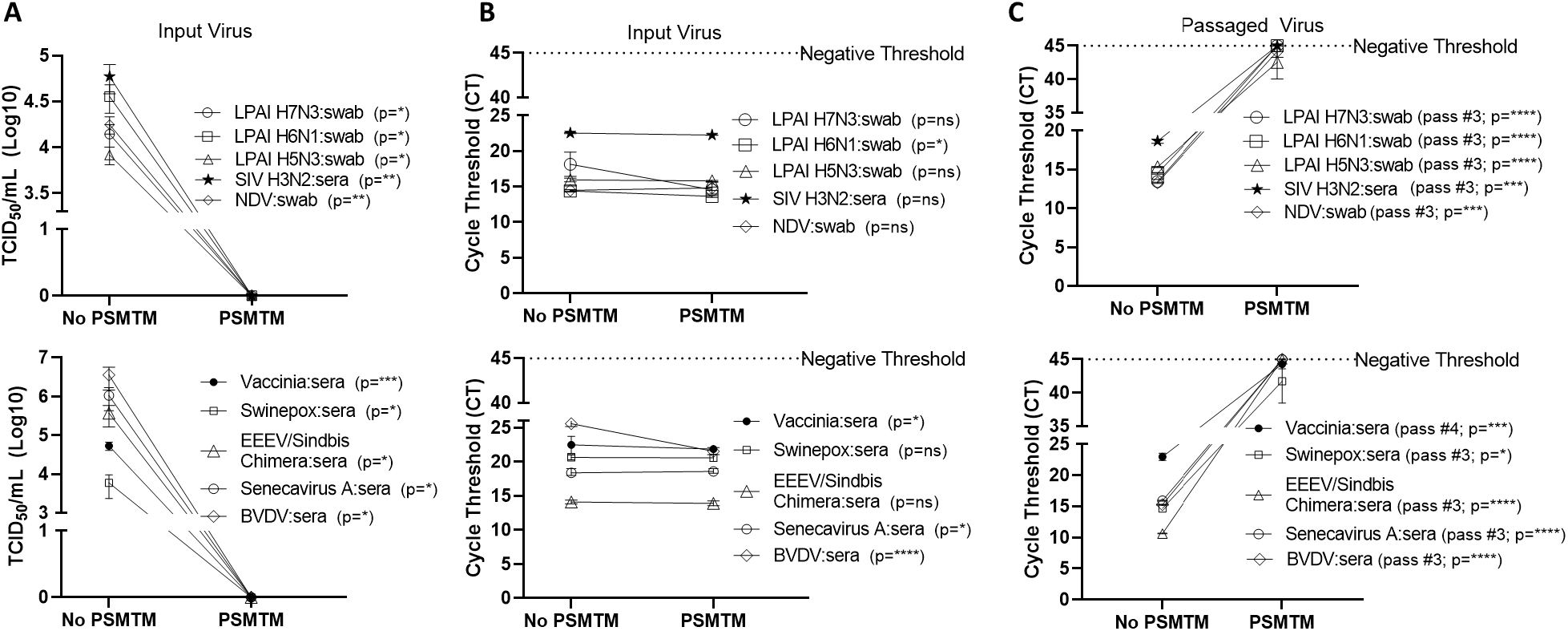
Presence of biological material does not affect effectiveness of PSMTM on inactivation or nucleic acid detection of diverse viruses. Viruses were combined with species-appropriate sera or cloacal swab as described in methods. A) Virus recovery after PSMTM inactivation or PBS no-inactivation control. Virus recovery was assessed by TCID_50_ titer using the cell line described in Table 1. B) Nucleic acid cycle threshold (CT) detection after PSMTM inactivation or PBS no-inactivation control. C) Serial passage endpoint nucleic acid cycle threshold (CT) detection after PSMTM inactivation or PBS no-inactivation. Serial passage was completed using the cell line in Table 1 and description outlined in methods. Nucleic acid was detected according to the reference in Table 1 and description in methods. PSMTM-treated viruses were compared to corresponding no-PSMTM control and inactivation was assessed using manufacturer recommended conditions. Significance was determined using the Student’s t-test. * *p*<0.05; ** *p*<0.01; *** *p*<0.001; **** *p*<0.0001; ns, not significant. Error bars represent standard error of the mean (SEM) of triplicate experiments.

### Maintenance of appropriate PSMTM ratio is required for effective virus inactivation

Collection of diagnostic samples may not always achieve exact 3 PSMTM: 1 virus inactivation ratio recommended by the manufacturer. To assess the safety of PSMTM inactivation use if the recommended ratio is not maintained, we evaluated virus infectivity and serial passage of virus treated with lesser amounts of PSMTM. EEEV/sindbis virus chimera was evaluated due to high titer growth. Results show that 1 PSMTM: 1 virus ratio completely inactivated virus infectivity >6-log and virus serial passage reached negative threshold by passage endpoint, whereas control-treated virus did not (Fig. 4A). However, although 1 PSMTM: 3 virus ratio inactivated virus infectivity >6-log, virus serial passage results revealed that virus was not completely inactivated where PSMTM-treated virus CT values were comparable to control-treated virus CT values by passage endpoint (Fig. 4B). Results of 1 PSMTM: 10 virus and 1 PSMTM: 100 virus ratio alterations further showed that virus was not inactivated as measured by infectivity and serial passage assays (Fig. 4C-D). These results indicate that minimal variations in the PSMTM inactivation ratio recommended by the manufacturer may still completely inactivate virus. However, caution should be used during sample collection to maintain the appropriate ratio to ensure complete inactivation.

**Figure 4:**
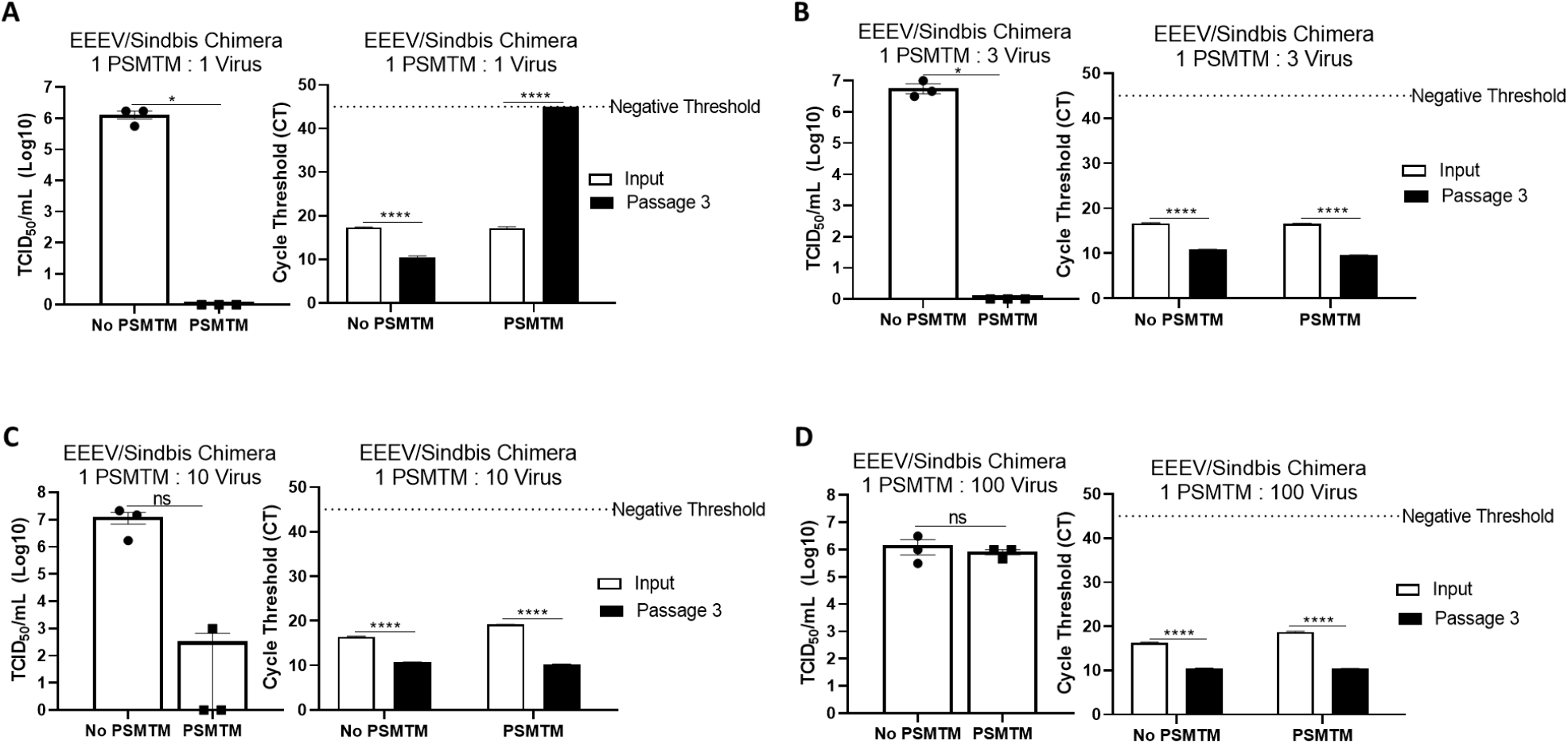
PSMTM to virus ratio determines effectiveness of inactivation. Virus recovery and nucleic acid detection were determined after PSMTM inactivation or PBS no-inactivation control utilizing various treatment to virus ratio conditions A) 1:1 ratio; B) 1:3 ratio; C) 1:10 ratio; D) 1:100 ratio. Virus recovery was assessed by TCID_50_ titer using the cell line described in Table 1. Nucleic acid was detected according to the reference in Table 1 and description in methods. PSMTM-treated viruses were compared to corresponding no-PSMTM control. Significance was determined using the Student’s t-test. * *p*<0.05; ** *p*<0.01; *** *p*<0.001; **** *p*<0.0001; ns, not significant. Error bars represent standard error of the mean (SEM) of triplicate experiments.

### PSMTM may modestly enhance stability of viral nucleic acid content required for PCR testing

Shipment of samples may take multiple days to reach laboratories for diagnostic testing, during which time nucleic acid may deteriorate or environmental conditions such as evaporation may influence inactivation efficacy [35]. We evaluated the effect of PSMTM on inactivation stability and virus nucleic acid detection over time. LPAI H7N3 and vaccinia virus were evaluated due to moderate and high stability, respectively, of infectivity over time at ambient temperature (Fig. 1A; 1F). PSMTM-treated or control-treated viruses were deposited into screw-cap tubes to mimic sample collection tubes and collected at the indicated timepoint. Virus infectivity for each timepoint was compared to timepoint 0 of each respective treatment.

Infectivity results show that PSMTM-treated viruses were completely inactivated at all timepoints compared to control-treated viruses (Fig. 5A; 5C). These data indicate that PSMTM inactivation is complete and virus infectivity is not recovered over time. Control-treated virus infectivity results reveal similar trends to our natural decay findings (Fig 1A; 1F), where LPAI H7N3 was moderately stable until complete loss of infectivity by day 14 (Fig. 5A). However, LPAI H7N3 nucleic acid was detected over 21 days despite loss of infectivity (Fig. 5A-B). Comparison of PSMTM- and control-treated LPAI H7N3 viruses show comparable CT values at each timepoint with only moderate increase in CT stability (reduced CT value or increased nucleic acid) in PSMTM at later timepoints (Fig. 5B). Control-treated vaccinia virus was highly stable, and infectivity was only reduced <2-logs by 21 days at ambient temperature (Fig. 5C). Vaccinia virus nucleic acid was detected over 21 days, and CT values were comparable for PSMTM- and control-treated viruses with slight increase in CT stability (reduced CT value or increased nucleic acid) in PSMTM at later timepoints (Fig. 5D). These data indicate that PSMTM may enhance stability of viral nucleic acid at later timepoints, although the effect is modest.

**Figure 5:**
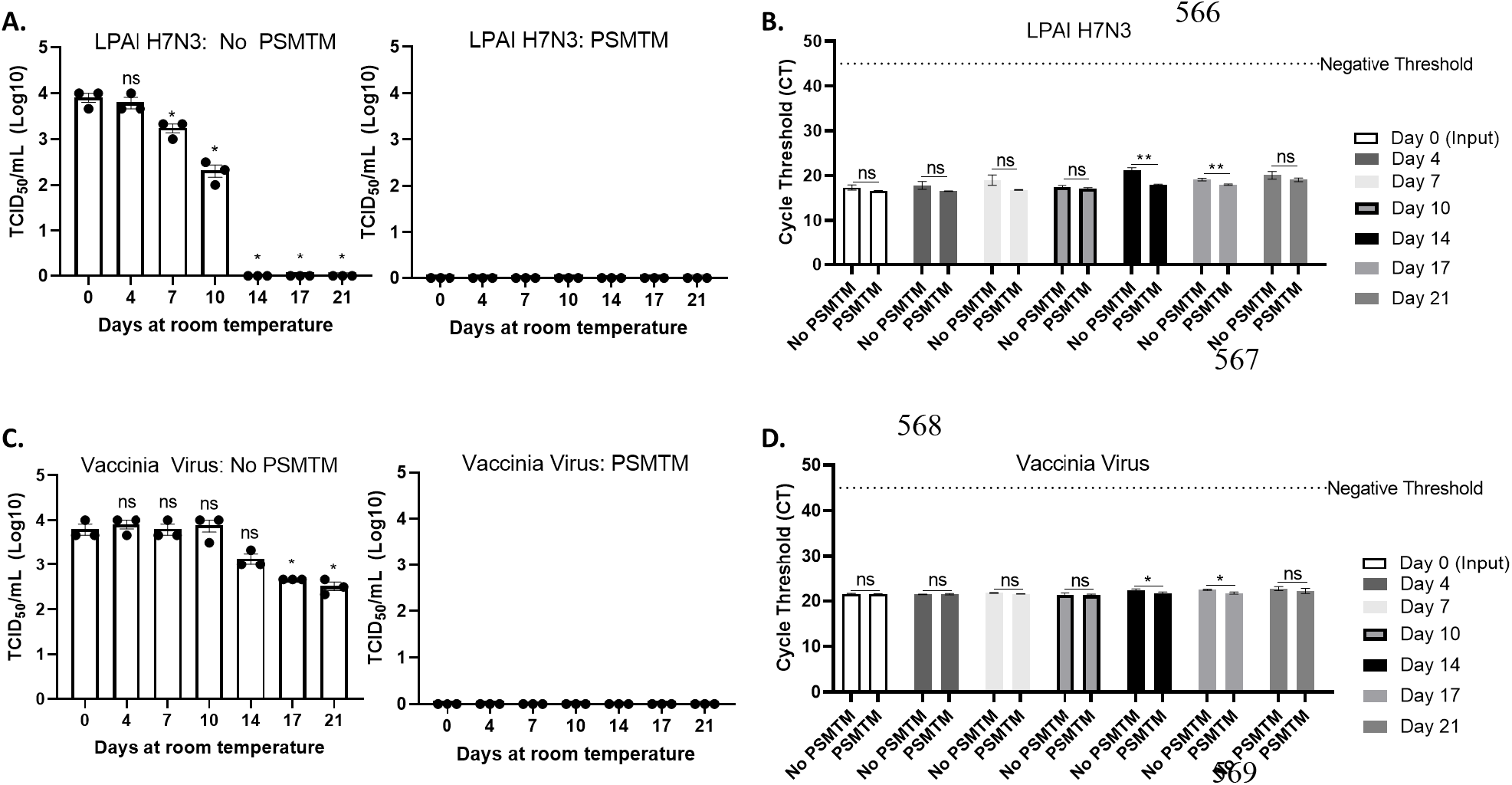
Virus inactivation and nucleic acid detection is stable over time in PSMTM. Virus recovery and nucleic acid detection over 21 days for viruses deposited in a plastic screw-cap tube at ambient temperature. Inactivation was assessed using manufacturer recommended conditions. A) Virus recovery was assessed by TCID_50_ for PBS no-inactivation control treated LPAI H7N3 (left) and PSMTM-treated LPAI H7N3 (right). B) Virus nucleic acid cycle threshold (CT) detection for PBS no-inactivation control and PSMTM-treated LPAI H7N3. C) Virus recovery was assessed by TCID_50_ for PBS no-inactivation control treated vaccinia virus (left) and PSMTM-treated vaccinia virus (right). D) Virus nucleic acid cycle threshold (CT) detection for PBS no-inactivation control and PSMTM-treated vaccinia virus. A,C) TCID_50_ titer was determined in the appropriate cell line as identified in Table 1. Each timepoint was compared to timepoint 0 for each respective treatment. B,D) Nucleic acid was detected according to the reference in Table 1 and description in methods. PSMTM-treated viruses were compared to corresponding no-PSMTM control. Significance was determined using the Student’s t-test. * *p*<0.05; ** *p*<0.01; ns, not significant. Error bars represent standard error of the mean (SEM) of triplicate experiments.

## Discussion

Highly pathogenic disease outbreaks, including the 2022 US HPAI outbreak and the COVID-19 pandemic, strain the already limited diagnostic capacity of high containment facilities. There is a need for a reagent that protects the nucleic acid required for diagnostic testing, while also inactivating pathogens to permit testing in lower containment facilities. PSMTM was previously utilized during the COVID-19 pandemic for diagnostic RT-PCR testing [24] and is authorized by the FDA for collection of IAV and *Mycobacterium tuberculosis* samples [17]. However, inactivation assessment of diverse viruses representing USDA Veterinary Services (VS) select agent viruses is needed due to the pathogenic potential and foreign animal disease designation of such viruses that may have devastating consequences on the food supply. In this study, we evaluated the inactivation of diverse animal virus surrogates, including four influenza A viruses, through titering and serial passage assays and assessed nucleic acid preservation through qRT-PCR.

The viruses used in this study showed reduced but detectable replication after several days at ambient temperature in liquid media (Fig. 1). Although conditions vary considerably between studies, these virus decay trends are in general agreement with those reported by others [36-43]. Therefore, an inactivation reagent utilized at the time of sample collection is ideal to alleviate concerns about transmission during shipping procedures and facilitate testing at lower containment facilities. All 10 viruses tested in this study, including four influenza A viruses of avian and swine origin, were completely inactivated (≥99.99% reduction) by PSMTM when utilized according to manufacturer recommendations (Fig. 2). Serial passage of PSMTM-treated viruses revealed no replicating virus (Fig. 2). Inactivation was not dependent on virus titer, as virus titers used in this study varied from 4-to 7-log/mL (Fig. 2-3). These titers are biologically relevant as animal model studies showed titers of ≤5log plaque forming units per mL (PFU/mL) for FMDV [44], 6-8log median hemadsorbing dose per mL (HAD_50_/mL) for ASF [45], 5-7log TCID_50_/mL for CSF [46], ≤7log PFU per gram tissue for EEEV [47] and IAV [48]; ≤7log and ≤8log TCID per gram tissue for NDV [49] and HPAI [50], respectively. Inactivation results were verified in the presence of biological material (virus-spiked sera or virus-spiked cloacal swabs) (Fig. 3) and are consistent with findings by others that showed chemical inactivation methods are not sensitive to the presence of biological material [33]. However, natural infection studies are needed to confirm that PSMTM inactivation is not sensitive to saturating concentrations of biological material.

Previous studies have shown that PSMTM inactivation is complete even with reduced contact time from the manufacturer recommended 60 minutes, where a contact time of as little as 2 minutes was identified as sufficient [51]. However, previous studies did not evaluate the importance of the volumetric ratio of inactivation media to virus-containing sample volume. Variation of the inactivation media ratio in our results showed that inactivation was incomplete with modest deviation from manufacturer recommended 3:1 ratio (Fig. 4). Titering and serial passage results revealed that inactivation was complete when the ratio was altered from the manufacturer recommended 3:1 to a 1:1 ratio; however, inactivation was incomplete with additional decrease of the inactivation media to virus sample media ratio (Fig. 4). Incomplete inactivation was not attributed to a significant increase in titer of virus sample media as the PBS without PSMTM inactivation control conditions were within 1-log for all ratio deviations (Fig. 4). While previous studies indicated that the contact time can be reduced without compromising inactivation, these results indicate that adherence to the manufacturer recommended volumetric ratio provides a reliable safety margin to ensure complete inactivation.

Importantly, nucleic acid CT values were within 1.5 CT value for all viruses in PSMTM and PBS no-inactivation control conditions indicating preservation of the nucleic acids despite inactivation of virus replication (Fig. 2-3). Other studies have shown CT values were comparable for detection of human respiratory viruses collected in PSMTM compared to other media [23, 52]. Together, the results of this study and those from previous studies by others show that PSMTM maintains the stability of both viral RNA and viral DNA for human and animal viruses despite the reduced environmental stability of viral RNA [23, 52, 53]. In addition, results from this study show inactivation and nucleic acid stability appear to be maintained over extended periods of time.

Incubation of LPAI H7N3 and vaccinia virus over 21 days revealed PSMTM inactivation was complete with no observation of virus replication despite comparable or modestly enhanced nucleic acid detection for PSMTM compared to PBS no-inactivation control (Fig. 5). The endpoint of this study was 21 days; however, other studies showed nucleic acid was detected from samples in PSMTM up to 196 days after collection [52].

In this study, PSMTM was completely effective at inactivation of all viruses tested. Our results are consistent with others that evaluated the use of PSMTM with human viruses [23, 52]. However, the viruses used here were surrogate viruses, and additional studies are needed with the target viruses represented in this study to validate inactivation with actual select agent viruses. This inactivation method may improve testing turn-around time during highly pathogenic disease outbreaks by de-restricting diagnostic testing. This method may also be used to ensure sample quality is maintained during normal shipping procedures. The results of this study may be applied to other pathogens that require molecular testing and may be sensitive to environmental conditions such as temperature and time to testing, and in scenarios where samples may be collected in field sites with limited resources.

## Author Contributions

JW, RL, SR, and KS contributed to study design and conception. JW and HH contributed essential study materials. JW, RS, NP contributed to data acquisition. JW performed statistical analysis. JW and RP wrote the first draft of the manuscript. All authors contributed to manuscript revision and read and approved the submitted version.

## Acknowledgments

We thank the Cytology group within the USDA National Veterinary Services Laboratories Diagnostic, Bioanalytical, and Reagents Laboratory Proficiency Testing and Reagents section for cell culture. We thank June Meichsner and Alethea Fry within USDA Center for Veterinary Biologics for providing primary CEF cells. We thank Denise Chapman in the USDA Animal Resources Unit for cloacal swab collection. We thank the USDA National Veterinary Services Laboratories Diagnostic Virology Laboratory for providing source viruses.

## Data Availability Statement

The contributions presented in the study are included in the article or supplementary material. Inquiries can be directed to the corresponding author.

## References

1. Belay, E.D., et al., Zoonotic Disease Programs for Enhancing Global Health Security. Emerg Infect Dis, 2017. 23(13): p. S65–70.

2. Powers, A.M., Resurgence of Interest in Eastern Equine Encephalitis Virus Vaccine Development. J Med Entomol, 2022. 59(1): p. 20–26.

3. Brown, V.R., et al., Risks of introduction and economic consequences associated with African swine fever, classical swine fever and foot-and-mouth disease: A review of the literature. Transbound Emerg Dis, 2021. 68(4): p. 1910–1965.

4. Thornber, P.M., R.J. Rubira, and D.K. Styles, Humane killing of animals for disease control purposes. Rev Sci Tech, 2014. 33(1): p. 303–10.

5. You, S., et al., African swine fever outbreaks in China led to gross domestic product and economic losses. Nature food, 2021. 2: p. 802–808.

6. Farahat, R.A., et al., The resurgence of Avian influenza and human infection: A brief outlook. New Microbes New Infect, 2023. 53: p. 101122.

7. Prempeh, H., R. Smith, and B. Muller, Foot and mouth disease: the human consequences. The health consequences are slight, the economic ones huge. BMJ, 2001. 322(7286): p. 565–6.

8. Corrin, T., et al., Eastern Equine Encephalitis Virus: A Scoping Review of the Global Evidence. Vector Borne Zoonotic Dis, 2021. 21(5): p. 305–320.

9. Ul-Rahman, A., et al., Zoonotic potential of Newcastle disease virus: Old and novel perspectives related to public health. Rev Med Virol, 2022. 32(1): p. e2246.

10. Horman, W.S.J., et al., The Drivers of Pathology in Zoonotic Avian Influenza: The Interplay Between Host and Pathogen. Front Immunol, 2018. 9: p. 1812.

11. Charlier, J., et al., Disease control tools to secure animal and public health in a densely populated world. Lancet Planet Health, 2022. 6(10): p. e812–e824.

12. Morse, S.A. and B.R. Quigley, Select agent regulations. Microbial Forensics, 2020: p. 425–39.

13. Waldrup, K.A. and T.H. Conger, Maintaining a vigilance for foreign animal diseases. Vet Clin North Am Food Anim Pract, 2002. 18(3): p. 379–87, v.

14. Saini, V., et al., A Cold Chain-Independent Specimen Collection and Transport Medium Improves Diagnostic Sensitivity and Minimizes Biosafety Challenges of COVID-19 Molecular Diagnosis. Microbiol Spectr, 2021. 9(3): p. e0110821.

15. Xia, H. and Z. Yuan, High-containment facilities and the role they play in global health security. Journal of biosafety and biosecurity, 2022. 2022 v.4 no.1(no. 1): p. pp. 1–4.

16. Enserink, M. and J. Kaiser, Biodefense research. Accidental anthrax shipment spurs debate over safety. Science, 2004. 304(5678): p. 1726–7.

17. Daum, L.T. and G.W. Fischer, Rapid and Safe Detection of SARS-CoV-2 and Influenza Virus RNA Using Onsite Quantitative PCR Diagnostic Testing From Clinical Specimens Collected in Molecular Transport Medium. J Appl Lab Med, 2021. 6(6): p. 1409–1416.

18. Clarke, C., et al., Novel molecular transport medium used in combination with Xpert MTB/RIF ultra provides rapid detection of Mycobacterium bovis in African buffaloes. Sci Rep, 2021. 11(1): p. 7061.

19. LHNVD. PrimeStore Molecular Transport Medium® (MTM). MSDS October 29, 2020 [cited 2023 January 31]; v2:[Available from: https://static1.squarespace.com/static/5f7dde76e3d6f427822285d4/t/5fbe6052fa04221c71f07ecd/1606312018859/US_SDS_Version02_Oct29_2020.pdf.

20. Honeywood, M.J., et al., Use of guanidine thiocyanate-based nucleic acid extraction buffers to inactivate poliovirus in potentially infectious materials. J Virol Methods, 2021. 297: p. 114262.

21. Wood, B.A., et al., Inactivation of foot-and-mouth disease virus A/IRN/8/2015 with commercially available lysis buffers. J Virol Methods, 2020. 278: p. 113835.

22. Lin, Q., et al., Sanitizing agents for virus inactivation and disinfection. View (Beijing), 2020: p. e16.

23. Daum, L.T., et al., A clinical specimen collection and transport medium for molecular diagnostic and genomic applications. Epidemiol Infect, 2011. 139(11): p. 1764–73.

24. Welch, S.R., et al., Analysis of Inactivation of SARS-CoV-2 by Specimen Transport Media, Nucleic Acid Extraction Reagents, Detergents, and Fixatives. Journal of Clinical Microbiology, 2020. 58(11).

25. Zhu, J.J., African Swine Fever Vaccinology: The Biological Challenges from Immunological Perspectives. Viruses, 2022. 14(9).

26. Ebling, R., et al., Virus viability in spiked swine bone marrow tissue during above-ground burial method and under in vitro conditions. Transbound Emerg Dis, 2022. 69(5): p. 2987–2995.

27. Rhee, C.H., M. Her, and W. Jeong, Modified Vaccinia Virus Ankara as a Potential Biosafety Level 2 Surrogate for African Swine Fever Virus in Disinfectant Efficacy Tests. Pathogens, 2022. 11(3).

28. Koonin, E.V. and N. Yutin, Origin and evolution of eukaryotic large nucleo-cytoplasmic DNA viruses. Intervirology, 2010. 53(5): p. 284–92.

29. Dee, S.A., et al., Survival of viral pathogens in animal feed ingredients under transboundary shipping models. PLoS One, 2018. 13(3): p. e0194509.

30. Mahy, B.W.J. and M.H.V. Van Regenmortel, Desk encyclopedia of animal and bacterial virology. 2010, Amsterdam ; London: Academic.

31. Wang, E., et al., Chimeric sindbis/eastern equine encephalitis vaccine candidates are highly attenuated and immunogenic in mice. Vaccine, 2007. 25(43): p. 7573–7581.

32. Weber, D.J., et al., The effect of blood on the antiviral activity of sodium hypochlorite, a phenolic, and a quaternary ammonium compound. Infect Control Hosp Epidemiol, 1999. 20(12): p. 821–7.

33. Welch, J.L., et al., Inactivation of Severe Acute Respiratory Coronavirus Virus 2 (SARS-CoV-2) and Diverse RNA and DNA Viruses on Three-Dimensionally Printed Surgical Mask Materials. Infect Control Hosp Epidemiol, 2021. 42(3): p. 253–260.

34. Al Bayat, S., et al., Can the cycle threshold (Ct) value of RT-PCR test for SARS CoV2 predict infectivity among close contacts? J Infect Public Health, 2021. 14(9): p. 1201–1205.

35. Morris, D.H., et al., Mechanistic theory predicts the effects of temperature and humidity on inactivation of SARS-CoV-2 and other enveloped viruses. Elife, 2021. 10.

36. Tuladhar, E., et al., Different virucidal activities of hyperbranched quaternary ammonium coatings on poliovirus and influenza virus. Appl Environ Microbiol, 2012. 78(7): p. 2456–8.

37. Osman, N., et al., Vaccine Quality Is a Key Factor to Determine Thermal Stability of Commercial Newcastle Disease (ND)Vaccines. Vaccines (Basel), 2021. 9(4).

38. Choo, J.J.Y., et al., Developing a Stabilizing Formulation of a Live Chimeric Dengue Virus Vaccine Dry Coated on a High-Density Microarray Patch. Vaccines (Basel), 2021. 9(11).

39. Newman, F.K., et al., Stability of undiluted and diluted vaccinia-virus vaccine, Dryvax. J Infect Dis, 2003. 187(8): p. 1319–22.

40. Caserta, L.C., et al., Stability of Senecavirus A in animal feed ingredients and infection following consumption of contaminated feed. Transboundary and Emerging Diseases, 2022. 69(1): p. 88–96.

41. Muhsen, M., et al., Biological properties of bovine viral diarrhea virus quasispecies detected in the RK13 cell line. Arch Virol, 2013. 158(4): p. 753–63.

42. Cardoso, N., et al., Bovine Viral Diarrhea Virus Infects Monocyte-Derived Bovine Dendritic Cells by an E2-Glycoprotein-Mediated Mechanism and Transiently Impairs Antigen Presentation. Viral Immunol, 2016. 29(7): p. 417–29.

43. Gibbs, P., Merck Veterinary Manual: Swinepox. 2021.

44. Colenutt, C., et al., Quantifying the Transmission of Foot-and-Mouth Disease Virus in Cattle via a Contaminated Environment. mBio, 2020. 11(4).

45. Guinat, C., et al., Dynamics of African swine fever virus shedding and excretion in domestic pigs infected by intramuscular inoculation and contact transmission. Vet Res, 2014. 45(1): p. 93.

46. Munoz-Gonzalez, S., et al., Postnatal persistent infection with classical Swine Fever virus and its immunological implications. PLoS One, 2015. 10(5): p. e0125692.

47. Honnold, S.P., et al., Eastern equine encephalitis virus in mice II: pathogenesis is dependent on route of exposure. Virol J, 2015. 12: p. 154.

48. Jacobsen, H., et al., Offspring born to influenza A virus infected pregnant mice have increased susceptibility to viral and bacterial infections in early life. Nat Commun, 2021. 12(1): p. 4957.

49. Hu, Z., et al., High levels of virus replication and an intense inflammatory response contribute to the severe pathology in lymphoid tissues caused by Newcastle disease virus genotype VIId. Arch Virol, 2015. 160(3): p. 639–48.

50. Kandeil, A., et al., Rapid evolution of A(H5N1) influenza viruses after intercontinental spread to North America. Nat Commun, 2023. 14(1): p. 3082.

51. van Bockel, D., et al., Evaluation of Commercially Available Viral Transport Medium (VTM) for SARS-CoV-2 Inactivation and Use in Point-of-Care (POC) Testing. Viruses, 2020. 12(11).

52. Schlaudecker, E.P., et al., Comparison of a new transport medium with universal transport medium at a tropical field site. Diagn Microbiol Infect Dis, 2014. 80(2): p. 107–10.

53. Kockler, Z.W. and D.A. Gordenin, From RNA World to SARS-CoV-2: The Edited Story of RNA Viral Evolution. Cells, 2021. 10(6).

54. Baker, J.L. and B.M. Ward, Development and comparison of a quantitative TaqMan-MGB real-time PCR assay to three other methods of quantifying vaccinia virions. J Virol Methods, 2014. 196: p. 126–32.

55. Kaiser, F.K., et al., Swinepox Virus Strains Isolated from Domestic Pigs and Wild Boar in Germany Display Altered Coding Capacity in the Terminal Genome Region Encoding for Species-Specific Genes. Viruses, 2021. 13(10).

56. Baxi, M., et al., A one-step multiplex real-time RT-PCR for detection and typing of bovine viral diarrhea viruses. Vet Microbiol, 2006. 116(1-3): p. 37–44.

57. Richt, J.A., et al., Real-time reverse transcription-polymerase chain reaction assays for the detection and differentiation of North American swine influenza viruses. J Vet Diagn Invest, 2004. 16(5): p. 367–73.

58. Joshi, V.G., et al., Prevalence of Newcastle Disease Virus in Commercial and Backyard Poultry in Haryana, India. Front Vet Sci, 2021. 8: p. 725232.

59. Armstrong, P.M., N. Prince, and T.G. Andreadis, Development of a multi-target TaqMan assay to detect eastern equine encephalitis virus variants in mosquitoes. Vector Borne Zoonotic Dis, 2012. 12(10): p. 872–6.

60. Dall Agnol, A.M., et al., A TaqMan-based qRT-PCR assay for Senecavirus A detection in tissue samples of neonatal piglets. Mol Cell Probes, 2017. 33: p. 28–31.

