## Supplemental figures for "Inactivation of Highly Transmissible Livestock and Avian Viruses Including Influenza A and Newcastle Disease Virus for Molecular Diagnostics"

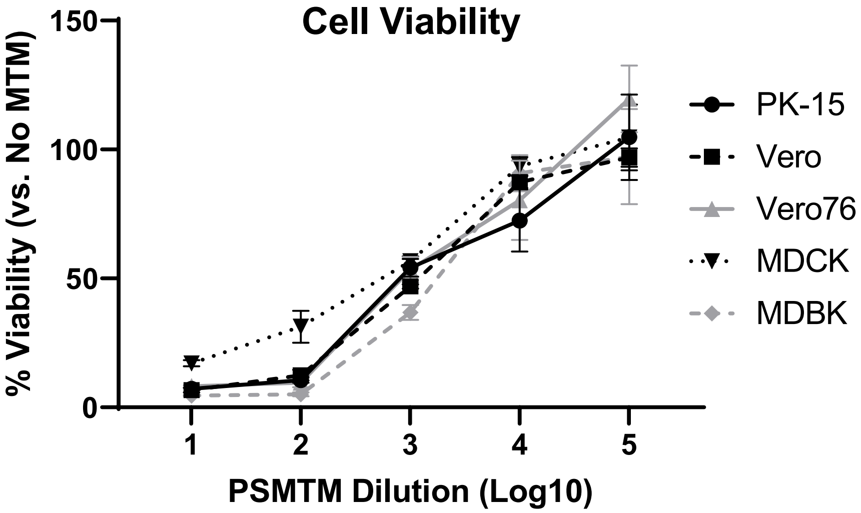


**Supplemental Figure 1: PSMTM is cytotoxic to a variety of immortalized cell lines.** PSMTM and PBS no-inactivation control were first diluted in virus propagation media (without virus) to manufacturer recommended conditions before additional 10-fold serial dilutions and deposited onto cells. Cell viability was determined by MTT assay for PK-15, Vero, Vero76, MDCK, and MDBK cell lines after incubation at 37°C for 48h. PSMTM cell viability was normalized to PBS no-inactivation control set at 100%. Error bars represent standard error of the mean (SEM) of triplicate experiments.

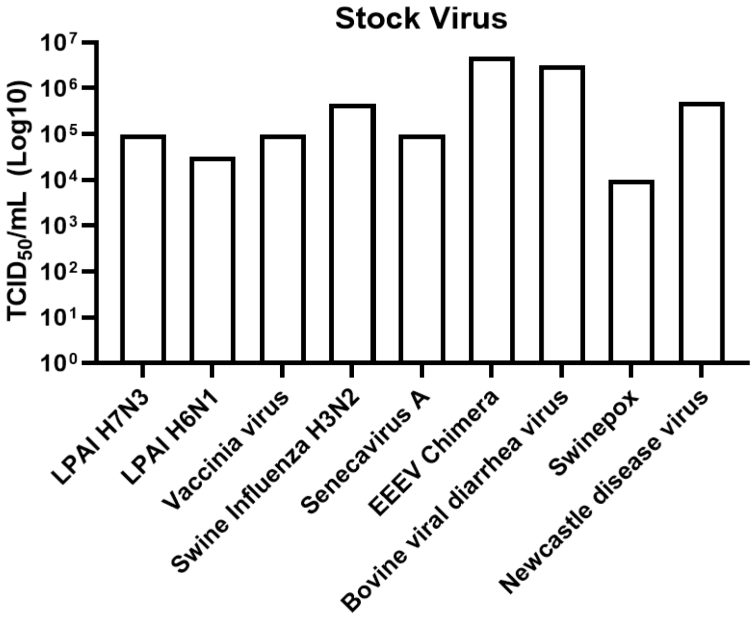


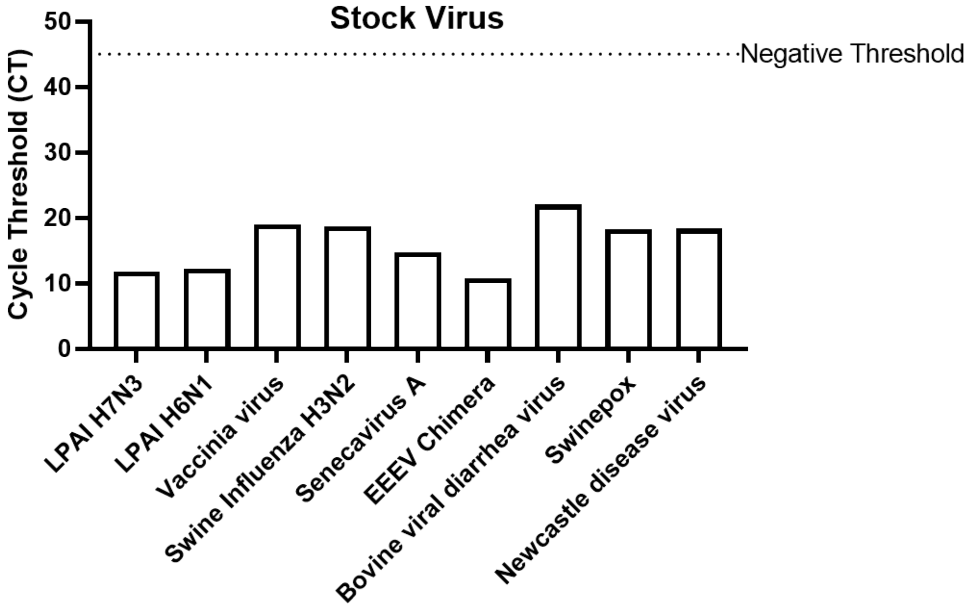


**Supplemental Figure 2: Stock virus $TCID_{50}$ and cycle threshold (CT) values reveal similar trends.** $TCID_{50}$ titers (top) for all viruses used in this study were determined in the appropriate cell line according to Table 1 and compared to corresponding nucleic acid CT values (bottom). Nucleic acid was detected according to the reference in Table 1 and description in methods.

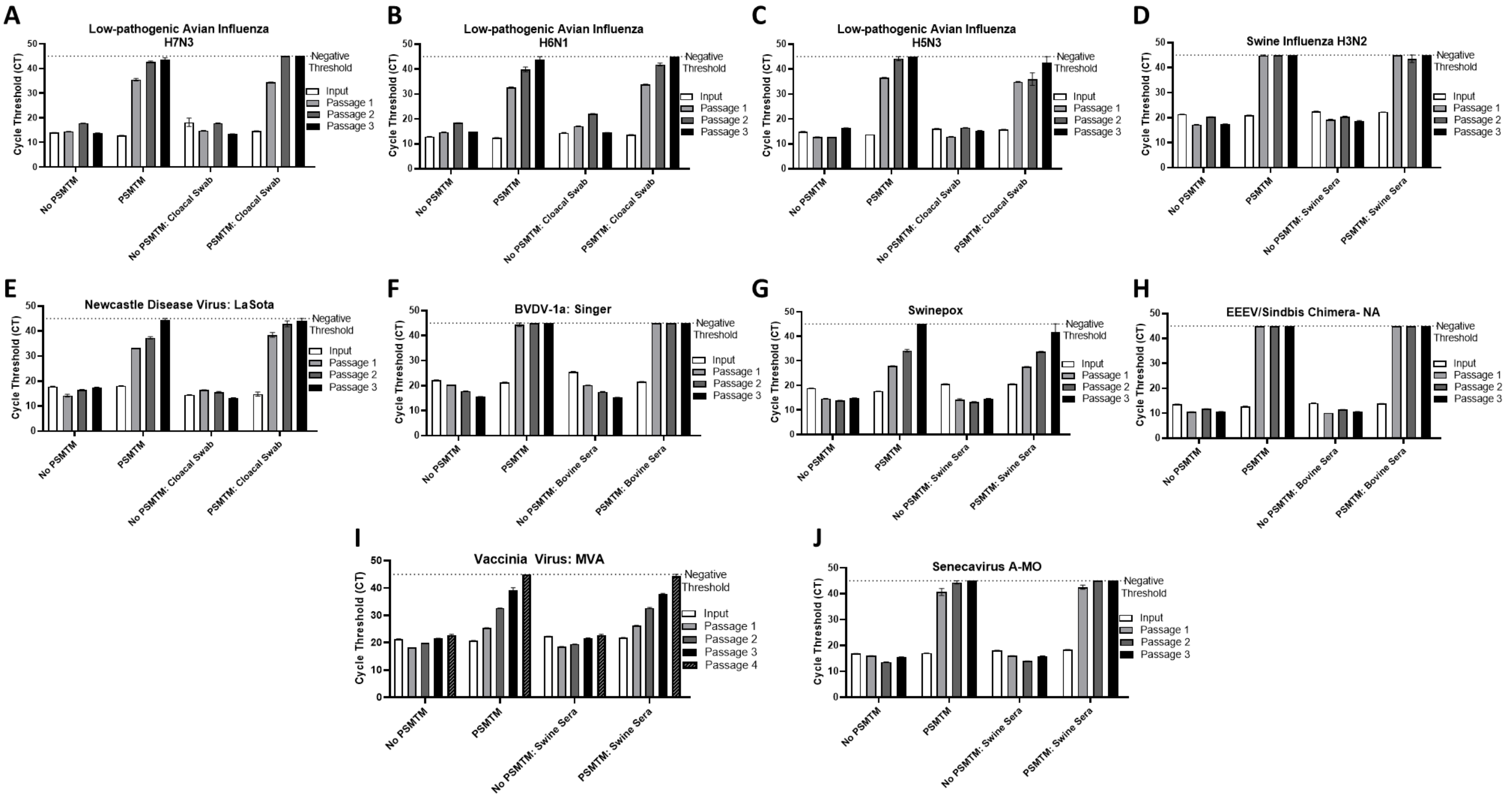


**Supplemental Figure 3: Serial passage of PSMTM-treated viruses shows decreased nucleic acid with each subsequent passage.** Nucleic acid cycle threshold (CT) detection for each serial passage after PSMTM inactivation or PBS no-inactivation control for A) LPAI H7N3; B) LPAI H6N1; C) LPAI H5N3; D) SIV H3N2; E) NDV; F) BVDV; G) Swinepox; H) EEEV/Sindbis chimera; I) vaccinia; and J) Senecavirus A. Inactivation was assessed using manufacturer recommended conditions. Serial passage was completed using the appropriate cell line in Table 1 and description outlined in methods. Nucleic acid was detected according to the reference in Table 1 and description in methods. Error bars represent standard error the mean (SEM) of triplicate experiments.

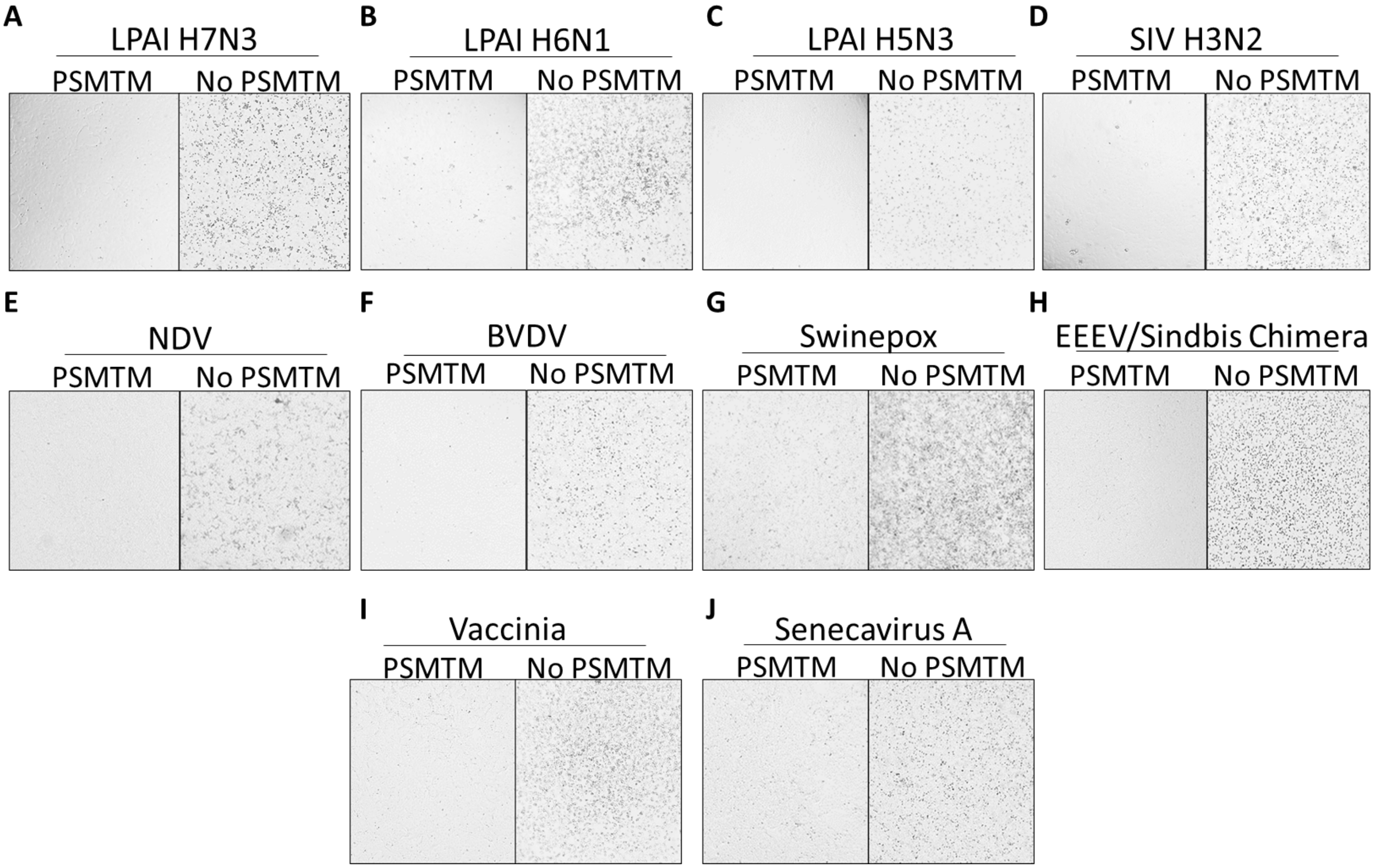


**Supplemental Figure 4: Cytopathic effect (CPE) visualization after serial passage of PSMTM-treated viruses shows no replicating virus.** CPE after three serial passages for PSMTM inactivation or PBS no-inactivation control for A) LPAI H7N3; B) LPAI H6N1; C) LPAI H5N3; D) SIV H3N2; E) NDV; F) BVDV; G) Swinepox; H) EEEV/Sindbis chimera; I) vaccinia; and J) Senecavirus A. Inactivation was assessed using manufacturer recommended conditions. Serial passage was completed using the appropriate cell line in Table 1 and description outlined in methods. Shown are one representative images from triplicate experiments at 4x magnification.
